# A decision-time account of individual variability in context-dependent orientation estimation

**DOI:** 10.1101/853754

**Authors:** Ron Dekel, Dov Sagi

**Affiliations:** Department of Neurobiology, The Weizmann Institute of Science, Rehovot 7610001, Israel

## Abstract

Following exposure to an oriented stimulus, the perceived orientation is slightly shifted, a phenomenon termed the tilt aftereffect (TAE). This estimation bias, as well as other context-dependent biases, is speculated to reflect statistical mechanisms of inference that optimize visual processing. Importantly, although measured biases are extremely robust in the population, the magnitude of individual bias can be extremely variable. For example, measuring different individuals may result in TAE magnitudes that differ by a factor of 5. Such findings appear to challenge the accounts of bias in terms of learned statistics: is inference so different across individuals? Here, we found that a strong correlation exists between reaction time and TAE, with slower individuals having much less TAE. In the tilt illusion, the spatial analogue of the TAE, we found a similar, though weaker, correlation. These findings can be explained by a theory predicting that bias, caused by a change in the initial conditions of evidence accumulation (e.g., prior), decreases with decision time (Dekel & Sagi, 2019b). We contend that the context-dependence of visual processing is more homogeneous in the population than was previously thought, with the measured variability of perceptual bias explained, at least in part, by the flexibility of decision-making. Homogeneity in processing might reflect the similarity of the learned statistics.

**Highlights:**

- The tilt aftereffect (TAE) exhibits large individual differences.
- Reduced TAE magnitudes are found in slower individuals.
- Reduced TAE in slower decisions can be explained by the reduced influence of prior.
- Therefore, individual variability can reflect decision making flexibility.

## Introduction

Visual context, from space and time, is known to influence the processing of visual information. Using basic visual properties, such as orientation, motion, and color, clear behavioral and electrophysiological effects have been identified (Clifford & Rhodes, 2005; Clifford et al., 2007; Gibson, 1937; Gibson & Radner, 1937; Kohn, 2007; Lamme & Roelfsema, 2000; Webster, 2011, 2015). With orientation features, the contextual orientation is known to lead to a perceptually salient shift in the perceived orientation (Gibson, 1937; Gibson & Radner, 1937; Schwartz, Hsu, & Dayan, 2007). When the context surrounds a target, this phenomenon is referred to as the tilt illusion (TI, Fig. 1A, Clifford, 2014; Gibson, 1937), and when the context precedes a target in time, it is referred to as the tilt aftereffect (TAE, Fig. 1B, Gibson & Radner, 1937; Webster, 2015). In both space and time (Schwartz et al., 2007), a contextual orientation of 20° clockwise to vertical leads to a *counterclockwise* shift in the estimated orientation, by a few degrees (see Fig. 1).

**Fig. 1.**
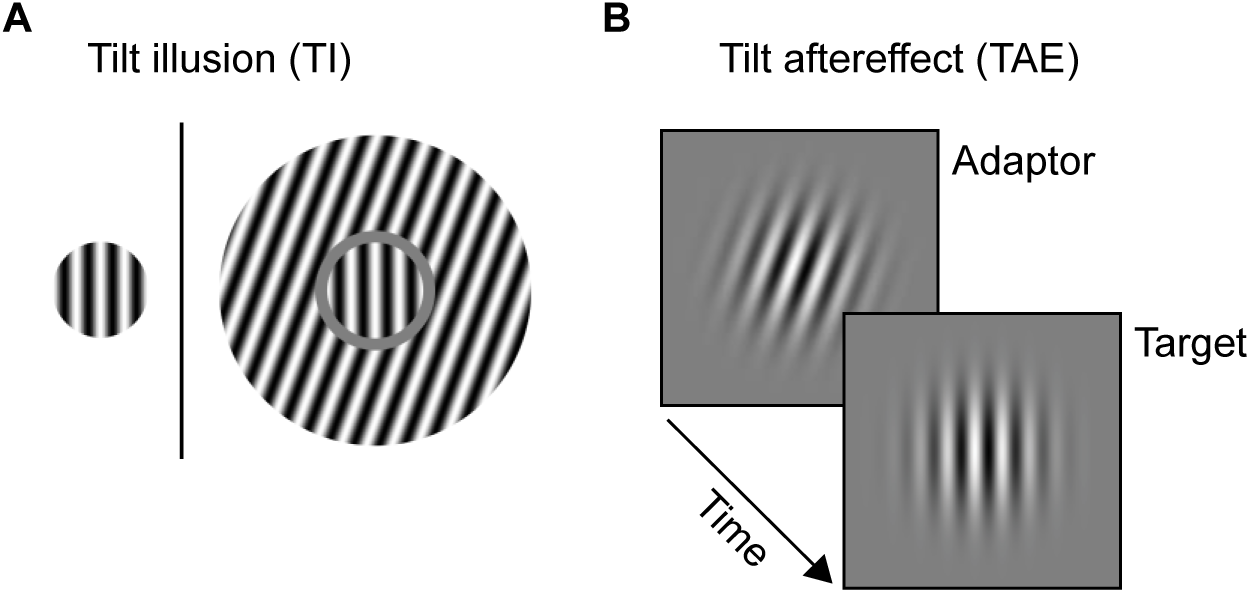
Tilt illusion and tilt aftereffect. (**A**) In the tilt illusion (TI), an oriented surround leads to a shift in perceived orientation. Right: the surrounding annulus, oriented 20° clockwise to vertical, leads to a *counterclockwise* shift in the perceived orientation of the center circle (the target). Left: the target without surround, was provided as a reference for the reader and was not used in the experiments. (**B**) In the tilt aftereffect (TAE), exposure to an oriented adaptor (e.g., +20°) leads to a shift in the perceived orientation of a subsequently viewed target (in the same direction as in TI). Figure reproduced from (Dekel & Sagi, 2019b).

Extensive theoretical work has been done to better understand such context effects. Generally, context-dependent changes in visual processing are thought to be functionally useful, despite some debate regarding details (Clifford, 2014; Kohn, 2007; Snow, Coen-Cagli, & Schwartz, 2017; Solomon & Kohn, 2014; Webster, 2011). Possible benefits include (a) self-calibration, constancy, or correction of a reference “norm” (Andrews, 1964; Day, 1972; Dekel & Sagi, 2019a; Gibson & Radner, 1937; Webster, 2011), (b) optimization of the neural code, such as improved gain of computational units, improved coding sensitivity to likely events, or decorrelation to remove coding redundancies (Benucci, Saleem, & Carandini, 2013; Coen-Cagli, Kohn, & Schwartz, 2015; Pinchuk-Yacobi & Sagi, 2019; Snow et al., 2017; Wei & Stocker, 2017), and (c) enhanced attentional selection of novel or surprising events (such events are presumably more likely to be important and hence deserve more attention). However, these and other alternatives are not necessarily mutually exclusive (e.g., orientation biases may reflect both self-calibration and decorrelation, Clifford, Wenderoth, & Spehar, 2000), and are not necessarily dependent on the neural implementation (e.g., divisive normalization may underlie both code optimization and attentional selection, Carandini & Heeger, 2012). However, two general observations can be made: First, theories seem to differ based on the speculated purpose (Press, Kok, & Yon, 2019), making perception more veridical (e.g., self-calibration), or less veridical but better at a given task (e.g., code optimization). Second, theories differ in how the effect is thought to depend on the computational constraints of the system. That is, if the system were to have better computational abilities (e.g., more neurons, more neural bandwidth), would context-dependent biases be less pronounced? For example, if biases reflect calibration based on the “true” white (white balance) or the “true” vertical, then it seems reasonable to assume that the biases are *not* dependent on computational constraints, and rather, are determined by a system-independent inference process (using stimulation statistics, such as the average color or the orientation modes). Alternatively, if biases reflect a tradeoff between computational constraints (such as limited bandwidth) and perceptual error, then we would expect less bias in a better system.

An interesting source of theory-diagnostic information can be obtained by considering individual differences in vision (Grzeczkowski, Clarke, Francis, Mast, & Herzog, 2017; Mollon, Bosten, Peterzell, & Webster, 2017). In the TI, as well as in other spatial-context-dependent biases, individuals measure large differences (by an order of magnitude), with strong test-retest reliability (Grzeczkowski et al., 2017; Song, Schwarzkopf, & Rees, 2013). These differences were found to be correlated with variability in orientation JND (just-noticeable-differences, showing an *R*^2^ value of ~60%) and were thought to reflect variability in the size of area V1 across individuals (Schwarzkopf, Song, & Rees, 2011; Song et al., 2013). These results seem consistent with an account of variability in terms of fixed neuronal constraints of low-level vision. In the TAE, direct investigation of individuality has, to the best of our knowledge, never been attempted, despite the large individuality evident in the literature (Gibson & Radner, 1937; Knapen, Rolfs, Wexler, & Cavanagh, 2010; Magnussen & Johnsen, 1986; Wolfe, 1984).

Importantly, we recently reported that context-dependent bias is much stronger in fast compared with slow reaction times (RT) of an individual (Dekel & Sagi, 2019b). This effect was largely independent of orientation sensitivity (i.e., JND). Moreover, we suggested that this within-observer variability in bias is explained by the theory that decision makers integrate evidence over time to reduce error, with an initial state of accumulation that is set by prior evidence favoring one decision outcome over others (Gold & Shadlen, 2007; Ratcliff, 1978; Ratcliff, Smith, Brown, & McKoon, 2016; Wald, 1945). In such models, bias, caused by the initial conditions, is expected to gradually decrease with decision time owing to noise accumulation, leading to a dramatic reduction of bias in slower decisions (Dekel & Sagi, 2019b; Mulder, Wagenmakers, Ratcliff, Boekel, & Forstmann, 2012; Ratcliff & McKoon, 2008; Summerfield & De Lange, 2014; White & Poldrack, 2014).

Here, we investigated individual differences in both TI and TAE, and considered their previously unexplored interaction with RT. Importantly, we found a strong negative correlation between TAE and RT, and a similar albeit statistically weaker correlation between TI and RT. These findings suggest that individual differences are based on RT, which complements current knowledge of differences in terms of orientation JND. This account is consistent with fixed low- and variable high-level visual processing.

## Methods

This study re-analyzed experimental data used in (Dekel & Sagi, 2019b).

### Observers

Twenty-nine observers (23 females, 6 males, aged 18-40) with normal or corrected-to-normal vision participated in the experiments. All observers were naïve to the purpose of the experiments, and provided their written informed consent. Most observers had prior experience of participation in psychophysical experiments. The work was carried out in accordance with the Code of Ethics of the World Medical Association (Declaration of Helsinki) and was approved by the Institutional Review Board (IRB) of the Weizmann Institute of Science.

### Apparatus

The stimuli were presented on a 22” HP p1230 monitor operating at 85Hz with a resolution of 1600×1200 that was gamma-corrected (linearized). The mean luminance of the display was 26 cd⋅m^−2^ (TAE experiments) or 49 cd⋅m^−2^ (TI experiments) in an otherwise dark environment. The monitor was viewed at a distance of 100 cm.

### Stimuli and tasks

All stimuli were presented using dedicated software on a uniform gray background. To begin stimulus presentation in a trial, observers fixated on the center of the display and pressed the spacebar (self-initiated trials). Responses were provided using the left and right arrow keys.

#### TAE experiments

The following presentation sequence was used (Fig. 1B): a blank screen (600 ms presentation), a Gabor “adaptor” (i.e., context, oriented −20° or +20° to vertical, 50 ms), a blank screen (600 ms), and a near-vertical Gabor “target” (50 ms). Observers were instructed to inspect the adaptor and target presentations, and then to report whether the orientation of the target was clockwise or counter-clockwise to vertical (no feedback). Gabor patches were 50% Michelson contrast, with a Gaussian envelope of σ = 0.42° and a sine wavelength of λ = 0.3° having a random phase. In the periphery experiments, adaptors and targets were presented at either left or right of the fixation (at ±1.8° eccentricity), the target was presented either at the same side as the adaptor (retinotopic) or at the opposite side (non-retinotopic), randomly, and targets were oriented from −12° to +12° (in steps of 2°). In the fixation experiment, adaptors and targets were presented at the fixated center of the display, and targets were oriented −9° to +9° (in steps of 1°). Four peripheral crosses co-appeared with the target to improve the discrimination between adaptor and target.

#### TI experiments

Stimuli (Fig. 1A right) consisted of a sine-wave circular “target” (radius of 0.6°) and a sine-wave annulus “surround” (width of 1.2°, and a gap of 0.15° from the central circle). Targets were oriented from −9° to +9° in steps of 1°, with λ = 0.3° and a random phase. Surrounding annuli were oriented −20° or +20°, with λ = 0.3° and a random phase. The contrast of the stimuli was 100%. Observers were instructed to inspect the target, and to report its orientation as clockwise or counter-clockwise to vertical (no feedback). The target+surround stimuli were presented starting from 350 ms after the trial initiation (“no jitter” experiment), or starting from 450 ms ± up to 100 ms (“onset jitter” experiment), for a duration of 200 ms.

### Procedure

In all conditions, each daily session was preceded by a brief practice block with easy stimuli (this practice was repeated until close-to-perfect accuracy was achieved).

#### TAE experiments

Sessions consisted of blocks lasting ~5 minutes, each with 125 trials. Blocks were separated by 2-minute breaks of blank screen-free viewing. In the periphery experiments, observers (*N* = 14) performed 3-8 daily sessions, each with five blocks. In the fixation experiment, observers (*N* = 12) performed a single session with six blocks.

#### TI experiments

Sessions consisted of blocks of 190 trials (lasting ~5 minutes), separated by 2-minute breaks of blank screen-free viewing. Observers (*N* = 10 for the “no jitter” experiment, and *N* = 10 for the “onset jitter” experiment) performed a single session with five blocks.

Note that the number of observers in the main experiment in this work (TAE in the periphery*, N* = 14) was relatively smaller than typically used to study individual differences. However, the number of repetitions per observer was relatively large (3-8 daily sessions of 625 trials), permitting higher precision in the individual measurement. All experiments in this study, with the exception of the TI “no jitter” experiment, were previously analyzed for the within-individual effects of RT (Dekel & Sagi, 2019b).

### Modeling

We consider two simple decision models that rely on accumulated evidence to make decisions, and try to approximate how model parameters relate to the behavioral measures of bias, sensitivity, and decision time. The models assume a temporal integration of evidence, at a fixed rate, with an initial state of accumulation that is set by prior evidence favoring (biasing) one decision outcome over others (Ratcliff, 1978; Ratcliff et al., 2016; Summerfield & De Lange, 2014). The integration of evidence is either stopped when a decision bound is reached (bounded model), or not (unbounded model), as detailed below.

In the analysis, we rely on the following definitions: First, Φ^−1^(⋅) is the inverse cumulative standard normal distribution function. In addition, Bias_crit_shift_ is the bias (e.g., TAE) measured by the shift of an internal criterion between the contexts (Dekel & Sagi, 2019; Green & Swets, 1966):

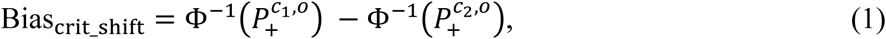

where 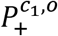 is the probability of clockwise answers for target orientation *o* under context *c*_1_ (e.g., a vertical target and a −20° adaptor exposure), and 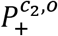 is the same for context *c*_2_ (e.g., the same target and a +20° adaptor exposure).

Finally, the orientation sensitivity, *d*’, is defined as (Green & Swets, 1966):

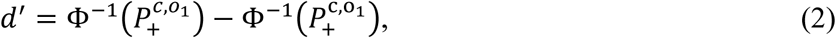

where 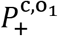 is the probability of clockwise answers for target orientation *o*_1_ under a given context *c* (e.g., a −0.5° target and a −20° adaptor exposure), and 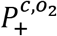 is the same for target orientation *o*_2_ (e.g., a +0.5° target and the same adaptor).

Based on standard signal detection theory modeling (SDT, Green & Swets, 1966):

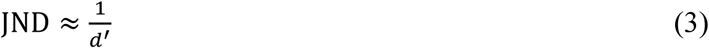

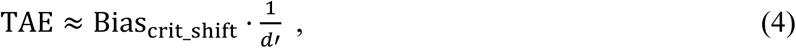

where JND is the just noticeable difference (the slope of the psychometric function), and TAE is bias measured in degrees (the shift of the psychometric function between the contexts).

#### Unbounded model

Here, we consider a simple case without decision bounds (an unbounded model). Thus, we assume some process of evidence accumulation that is equivalent to a simple random walk (a Wiener process) that starts from point *z* (which is a scalar) and gradually diverges due to stochastic diffusion (noise) and drift *v* (the delta of evidence being accumulated at each time point). Intuitively, we expect the influence of the starting point of the random walk to diminish at a rate proportional to the square-root of the time. More formally, the probability density of the random walk at decision time *t* is a normal distribution with mean *z* + *t* ⋅ *v* and variance *t*, namely, 𝒩(*z* + *t* ⋅ *v*, *t*). Therefore,

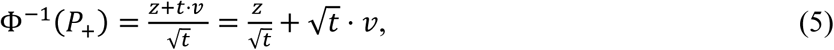

where *P*_+_ is the probability of the process being positive at time *t*. Then, the bias in the internal criterion (Bias_crit_shift_), given by the change in Φ^−1^(*P*_+_) due to the context (+*z* compared with −*z*, see Eq. (1)), is:

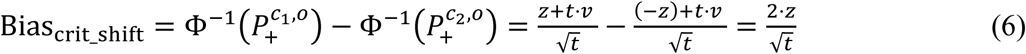

And, similarly, the sensitivity (*d*’), given by the change in Φ^−1^(*P*_+_) due to a change in the stimulus (+*v* compared with −*v*, see Eq. (2)), is:

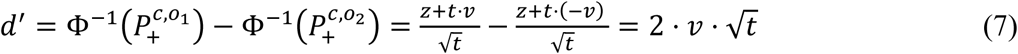

We now consider how the different behavioral measures are predicted to change, depending on the decision time *t*. Using Eqs. (3)-(7), and assuming that the target orientation is linear in the *v* parameter, we obtain:

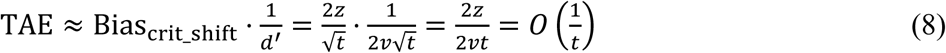

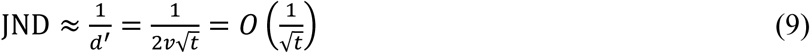

Based on Eqs. (8) and (9), the unbounded model makes the following predictions concerning changes in TAE and JND for a changed decision time:

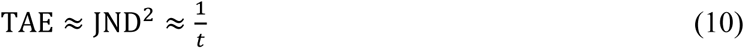

Note that the unbounded model does not explain the decision time itself, only how the decision time affects bias and sensitivity.

#### Bounded model (DDM)

An alternative approach of modeling decision processes in the brain, which also explains decision times, is to assume that there are decision bounds. When the accumulated evidence reaches a bound, the process is stopped and a decision is made. Here we considered the standard bounded drift diffusion model (DDM, Gold & Shadlen, 2007; Ratcliff & McKoon, 2008; Ratcliff & Smith, 2015; Ratcliff et al., 2016; see mathematical background at Luce, 1986; Shadlen, Hanks, Churchland, Kiani, & Yang, 2006). The DDM can be defined using four parameters: the drift rate (*v*), bound separation (*a*), starting point (*z*), and non-decision time (*t*_0_). In this description, the bounds are at 0 and *a*, and the process starts from the point *z*. We also define *b* as the distance of the starting point from the midpoint between the bounds: 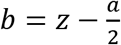.

Using Eq. (A.12) from (Palmer, Huk, & Shadlen, 2005) for the case *v* = 0, we get: 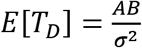 where *E*[*T*_*D*_] is the expected decision time (for both upper and lower bounds), *A* and *B* are the distances from the starting point to the upper and lower bounds, respectively, and *σ* is a scaling constant that is usually set to a fixed value of 1. Therefore, using *A* + *B* = *a* and 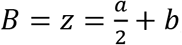, we get:

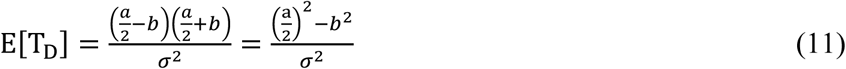

Moreover, using Eq. (A.6) from (Palmer et al., 2005) again for *v* = 0, we obtain 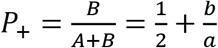, where *P*_+_ is the probability to reach the upper bound. Because 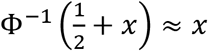 for small *x* values, we expect that when 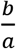 is not too large, it will approximately hold that 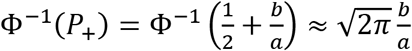. The bias, given by the change in Φ^−1^(*P*_+_) due to a change in context (+*b* compared with −*b*, see Eq. (1)), for the case where *v* = 0, is given by:

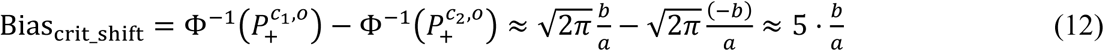

We consider now the task sensitivity. Using Eq. (A.13) from (Palmer et al., 2005) for the case of an unbiased starting point (*b* = 0), we obtain 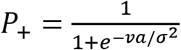. This is a logistic function, and thus, we can approximate it by 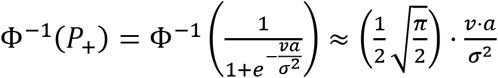. As such, the sensitivity (*d*’), given by the change in Φ^−1^(*P*_+_) due to a change in stimulus (Eq. (2)), when *b* = 0, is given by:

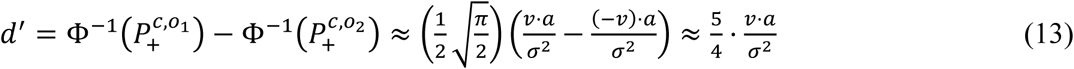

Although Eq. (12) was derived when there is a small 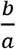 value, and Eq. (13) was derived when *b* = 0, we found the approximations to be reasonably robust in the relevant parameter range (Fig. 2). Based on the above approximations, we next determined how the different behavioral measures are predicted to change when only the bound separation (*a*) parameter is changed. Using Eqs. (3), (4), and (11)–(13), and also assuming that the target orientation is linear in the *v* parameter, we obtain:

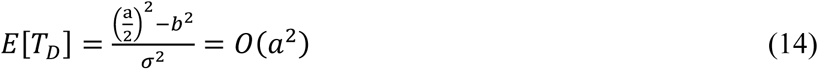

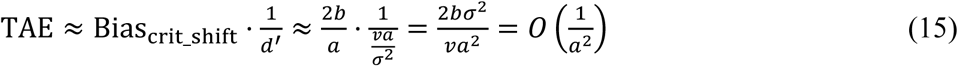

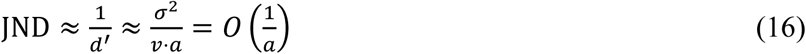

**Fig. 2.**
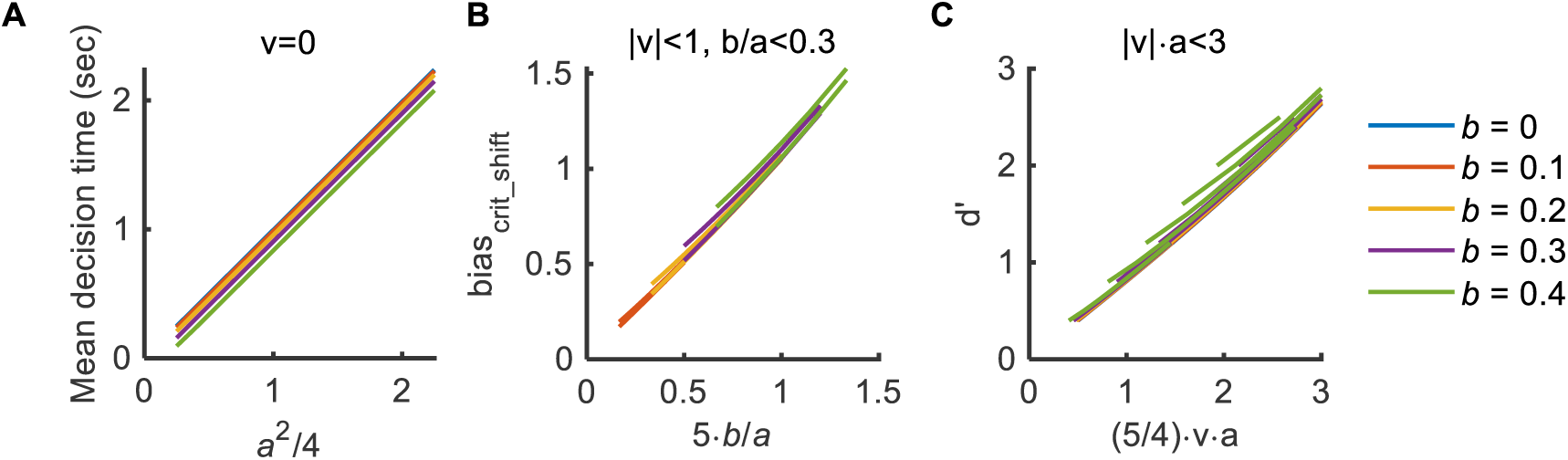
Robustness of Eqs. (11)-(13). Shown are true analytical DDM predictions, obtained using a computer analysis software (fast-dm, Voss & Voss, 2007), as a function of the derivations/approximations described in (**A**) Eq. (11), (**B**) Eq. (12), and (**C**) Eq. (13). In the figure, each plotted line was obtained by modifying the bound separation parameter (a). The different lines correspond to different values of the v (drift rate) and the b (the offset of the starting point from the mid-point) parameters, for the parameter range described in the title. Parameter values were further restricted to the behaviorally relevant ranges: 1 ≤ a ≤ 3, |v| ≤ 5. Panel (A) is for v = 0 because the relevant behavioral measure of decision time is RT for the vertical target orientation (RT_vert_). Overall, it can be seen that Eqs. (11)-(13) are practically linear with true DDM predictions.

Based on Eqs. (14)-(16), changing the bound separation (*a*) leads to approximately the following scaling:

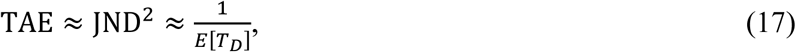

where *E*[*T*_D_] is the mean decision time.

In terms of the SPRT (Sequential Probability Ratio Test, Moran, 2015; Summerfield & De Lange, 2014; Wald, 1945), which is a standard Bayesian interpretation of the DDM, an increased bound separation corresponds to requiring lower type I and type II error rates, but starting from the same prior value. Hence, it seems reasonable to assume that the starting point remains the same when the bounds are changed.

#### Final remarks

As shown above, both models can predict the same dependence on decision time (Eqs. (10) and (17)). It is worthwhile to mention two important ways in which these models are over-simplified. First, both models assume zero noise at *t* = 0, which is clearly impossible in a biological system. Second, both models assume that the evidence is fixed in time; however, in many experiments (and here), the evidence is temporary (see the Discussion). Importantly, although modeling these constraints leads to a more complicated analysis, the TAE value is still expected to be reduced with decision time.

In addition, it seems interesting to extend the above analysis to take into account the possibility of individual differences in the rate of evidence accumulation (the drift rate, *v*). For the unbounded model, based on Eq. (9), we obtain 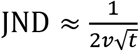; therefore 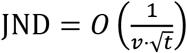. In addition, from Eq. (8), we obtain 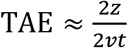; therefore 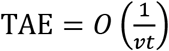. Combined, changing both the decision time and the drift rate leads

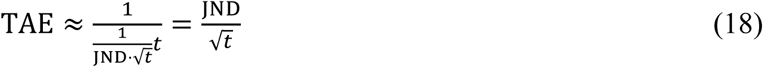

Consistent with Eq. (10). The same scaling is found in the DDM, when both the bound separation (*a*) and the drift rate are changed between observers. Finally, note that in all of the above analyses, we assumed that TAE only results from a change in the starting point of the process (bounded or unbounded). However, based on (Dekel & Sagi, 2019b), it is possible that the context leads to a change in both the starting point and the drift rate. In this sense, Eqs. (10) and (17) model the time-dependent component, with the total TAE having an additional time-independent, possibly additive, component.

### Analysis

#### Fitting the perceived orientation

The magnitude of TAE and TI was calculated based on the reported orientation (clockwise vs. counterclockwise) of the near-vertical targets (the Gabor patch for TAE, and the central sine-wave circle for TI). The perceived vertical orientation (PV) is the interpolated orientation having an equal probability for clockwise and counter-clockwise reports (50%) of the target. The PV was found separately for each context orientation (−20° and +20°). This was achieved by interpolation from a fit to a cumulative normal distribution (with lapse rates) of the psychometric function (the percent clockwise reports as a function of target orientation). Psignifit 3.0 software (Fründ, Haenel, & Wichmann, 2011) was used for fitting. Then, the TAE or TI magnitude was calculated as half the shift in PV between the two opposing adaptor or surround orientations (−20° vs. +20°), and, if relevant, averaged over the left and right target positions. We noted that an alternative measure of bias, Bias_crit_shift_, described above (Eq. (1)), can be defined from signal detection theory (Green & Swets, 1966). We found this alternative measure to be more convenient for RT-based modeling (Dekel & Sagi, 2019b); however, we preferred using the shift in the psychometric function because (i) Bias_crit_shift_ is less convenient in practical use because of its saturation with a large bias, and, at least here, loss of most of the collected data (i.e., using a single target orientation to calculate bias discards the data of the other target orientations); (ii) the shift in the psychometric function is expected to be more robustly correlated with RT, because, based on the modeling above, Bias_crit_shift_ is inversely proportional to the square-root of the time, whereas the shift in the psychometric function is inversely *linear* with time; (iii) last but not least, we preferred using a standard measure of TAE over a less standard one. Nevertheless, we verified that the main finding reported here, of a lower TAE in the slower observers, is found when measuring TAE using Bias_crit_shift_ (Eq. (1)) (data not shown).

To calculate JNDs (just noticeable differences), we used a multiple of the interpolated inverse slope of the psychometric function at the PV orientation. Specifically, we used the width of the interval over which the fitted cumulative Gaussian function rises from 0.25 to 0.75 (corresponding to 75% accuracy, or, put differently, to 1.35σ where σ is the fitted standard deviation). This interval was corrected so that the upper and lower lapse rates (as measured by the fit) do not affect the JND. The JNDs were calculated separately and then averaged over the −20° and +20° contexts, and, if relevant, over the left and right target positions.

#### Reaction times (RTs)

As a measure of RT for an observer, we used either the mean RT for all trials having all target orientations, or the mean RT only for trials having a vertical target orientation (RT_vert_). The first seems like a reasonable theory-agnostic measure, and the second is motivated by theory (see below and Dekel & Sagi, 2019). Importantly, the context leads to a shift in the psychometric function, which affects RTs (slowest RT at the *perceived*, i.e., the most difficult, target orientation, Ratcliff, 2014). Therefore, it is important to verify that using RT_vert_ does not introduce confounds for correlations with TAEs. Here, we verified this in two ways: first, by replicating the main findings using the interpolated RT at the PV orientation, which showed results almost identical to those reported. Second, conceptually, by measuring the size of the expected difference between RT_vert_ and the RT at the PV, which showed values around ~65 ms, and at most 220 ms, which is clearly negligible compared with the range of individual differences in RT_vert_ (see the Results). Where relevant, RTs were averaged over the left and right target positions.

#### Statistics

To obtain a measure for the statistical significance of individual differences in a factor, we considered repeated measurements over different days, fitting a linear mixed-effects model with one overall intercept term and also one term per observer, and reported the significance of an *F*-test of the null hypothesis that the coefficients of all observer terms are 0 (so only the overall intercept term remains). This analysis is only applicable for the periphery experiment, requiring multiple daily measurements. To obtain a measure for the co-variation of two factors, we used the Pearson correlation coefficient (*R*, or occasionally its square-root, *R*^2^, which measures shared variation). The statistical significance was assessed using the standard approach of applying a two-tailed *t*-test after transforming the data using Fisher’s *z*-transformation. An alternative approach of using a linear mixed-effects model shows the same results (with better significance), but it is only applicable for periphery data with multiple daily measurements. All statistical analyses were performed using MATLAB^®^ R2019b software. Equation (19) was fit with the “fitnlm” function, which finds the least-squares fit.

## Results

Using briefly presented Gabor patches (50 ms), we measured the shift in the estimated vertical orientation of a near-vertical Gabor (“target”), caused by previous exposure to a Gabor patch tilted −20° or +20° to vertical (“adaptor”, 600 ms ISI) (Fig. 1B). This experiment was performed using Gabor patches presented at the near-periphery (±1.8° eccentricity), permitting analysis based on the relative retinal positions of the adaptor and the target: the same position (“Retinotopic”) or contra-lateral positions (“Non-retinotopic”). The results for both retinotopic and non-retinotopic measurements showed a shift in the perceived orientation in the direction that is *away* from the adaptor orientation, a phenomenon known as the tilt aftereffect (TAE).

### Co-variation of RT and peripheral TAE

Importantly, TAE measured large individual variability (see the y-axis of Fig. 3), showing, in the retinotopic condition, magnitudes ranging from 0° to 2° (Mean ± *SD* of 1.04° ± 0.63°), and a very low measurement error (within-individual *SEM* over daily repetitions of ~0.2°). Statistically, the presence of an individual component in TAE was extremely significant (*p* = 1.9×10^−12^, *F*_(13,57)_ = 12.72, using a linear mixed-effects model, see the Methods). The non-retinotopic condition showed the same, albeit at weaker TAE magnitudes (Mean ± *SD* of 0.33° ± 0.36°, the individual component at *p* = 1.6×10^−5^, *F*_(13,57)_ = 4.77).

**Fig. 3.**
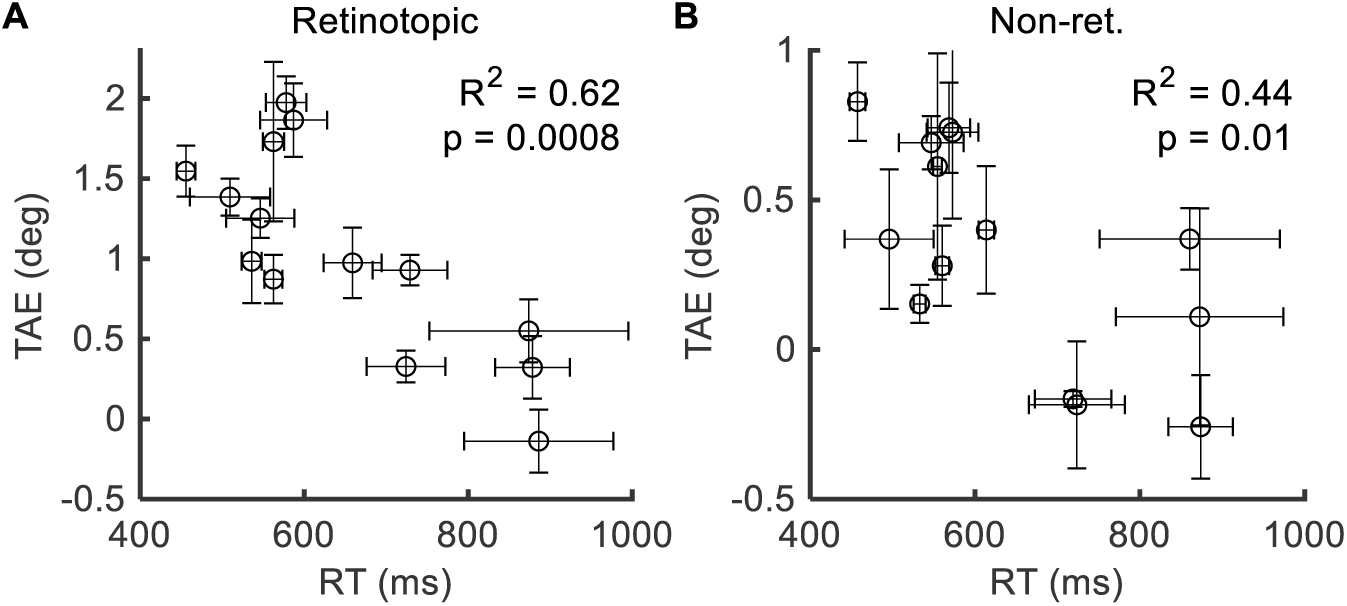
Co-variation of TAE and RT. TAE is shown as a function of the mean reaction time (RT), for different observers, in the (**A**) retinotopic and (**B**) non-retinotopic measurements. Error bars are SEM across daily repetitions. The results showed large individual differences in TAE that are largely accounted for by individual differences in RT.

Reaction time (RT), averaged across the easy and difficult target orientations, measured values ranging from about 500 to 1000 ms (see the x-axis of Fig. 3). Remarkably, the variability in TAE and RT was strongly and negatively correlated (Fig. 3), with the fast observers having much more TAE than the slow observers. Specifically, for the retinotopic measurement, we found TAEs of ~1.7° for observers with mean RTs of ~500 ms, and close to zero TAE for observers with RTs approaching 1000 ms (Fig. 3A) (*R*^2^ = 0.62, *p* = 0.0008, *t*_(12)_ = −4.47, two-tailed *t*-test following Fisher’s *z*-transformation, see the Methods). The non-retinotopic measurement revealed the same trend, albeit for weaker TAEs (Fig. 3B) (*R*^2^ = 0.44, *p* = 0.01, *t*_(12)_ = −3.05). Overall, variability in peripheral TAEs seems to be largely explained by RTs.

### Individual variability and decision times

To explain the co-variation of RT and TAE, we relied on the idea that decisions are made by an evidence-accumulation process with biased initial conditions, which leads to reduced TAE with decision time (see the Introduction) (Dekel & Sagi, 2019b). This is a general argument, and depending on modeling details, it can predict different rates of reduction in bias. Here, we considered the case where the rate of reduction in bias is inversely proportional to the decision time (i.e., RT minus non-decision time), as in Eqs. (10) and (17):

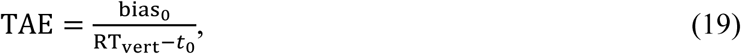

where bias_0_ is the initial bias at decision time zero, RT_vert_ is the RT for a vertical target, and *t*_0_ is the non-decision time (which reflects non-decision time components, such as the time it takes to press a response key, Ratcliff & McKoon, 2008). This equation reflects somewhat general assumptions that are broader than those of a single model. For example, Eq. (19) is predicted from an unbounded decision process (see Eq. (10)), and also from a bounded decision process with variability in the size of the separation between the bounds (see Eq. (17)). In addition, note that Eq. (19) might multiplicatively depend on the orientation sensitivity (if it is not fixed in the sampled population; see below, in the Discussion, and see Eq. (18)).

The measured RT_vert_ (the RT for the vertical target) exhibited a dramatic individual variation, with values ranging from 500 to almost 2000 ms (see the x-axis of Fig. 4). This range seems especially large when taking into account the non-decision time, *t*_0_, which, based on the RT distributions, we estimated to be about 350 ± 50 ms (Mean ± *SD* across observers, data not shown). Remarkably, RT_vert_ was strongly correlated with both retinotopic TAE (*R*^2^ = 0.74, *p* = 8×10^−5^, *t*_(12)_ = −5.85) and non-retinotopic TAE (*R*^2^ = 0.52, *p* = 0.004, *t*_(12)_ = −3.61). The correlations were stronger than those found with RT averaged over all target orientations, supporting the use of RT_vert_ to predict TAE. (Generally, the use of RT_vert_ was motivated by the idea that the relevant RT for bias is around the physical or the perceived vertical target orientation. We noted that the shifts in perceived orientation due to context led to a change in RT_vert_, but a small one, of ~65 ms, which is negligible compared with the range of the observed individual differences in RT_vert_, see the Methods.)

**Fig. 4.**
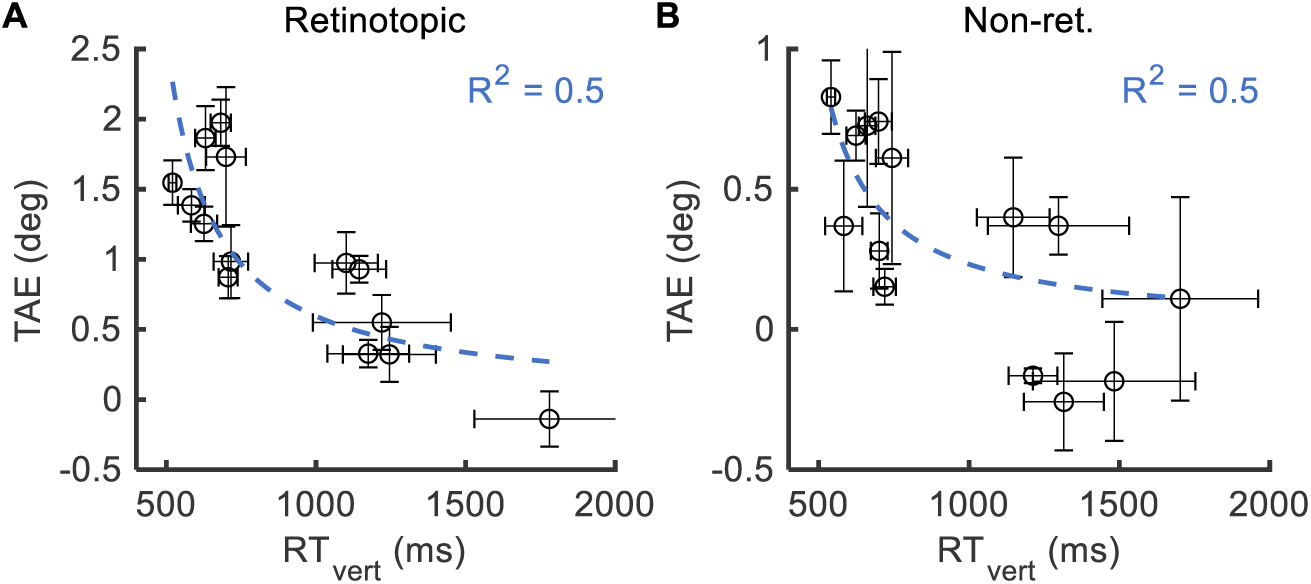
An RT account of the TAE. Shown is TAE as a function of the average RT for the vertical target orientation for different observers, in the (**A**) retinotopic and (**B**) non-retinotopic measurement. Blue dashed lines denote fits to Eq. (19) when setting t_0_ = 350 ms. Error bars are SEM across daily repetitions. Note that unlike Fig. 3, here the x-axis is the RT of the vertically oriented targets.

We next tried to account for individual differences using Eq. (19) (Fig. 4). We assumed that the initial bias (bias_0_) and the non-decision time (*t*_0_) are fixed in the population, and set *t*_0_ = 350 ms based on the RT distributions. Fitting bias_0_ to behavior showed that 50% of the variability in retinotopic and non-retinotopic TAEs can be explained by RT (*R*^2^ = 0.50, Fig. 4). Note that the predicted reduction by the fitted model is quite dramatic: from about 2° when RT = 500 ms, to about 0.5° when RT = 2000 ms in the retinotopic TAE (the dashed blue line in Fig. 4A). Fitting both bias_0_ and *t*_0_ to behavior showed similar results (retinotopic: *t*_0_ = 145 ms, *R*^2^ = 0.63; non-ret.: *t*_0_ = 368 ms, R^2^ = 0.5).

### Co-variation of RT and bias: the effects of experience

First, we restricted the analysis of peripheral TAE to the first experimental session, to minimize potential interaction with perceptual learning (Sagi, 2011). The results showed that the correlation between TAE and RT_vert_ is maintained, and that it is even stronger than when measurements are averaged across days (Fig. 5AC) (retinotopic: *R*^2^ = 0.68, *p* = 0.0005, *t*_(11)_ = −4.87, non-ret.: *R*^2^ = 0.5, *p* = 0.007 *t*_(11)_ = −3.33; one observer was not included in the analysis because of prior participation in a similar experiment). Similarly, fitting to Eq. (19) showed possibly better fits (with *t*_0_ = 350 ms, retinotopic: R^2^ = 0.69, non-ret.: R^2^ = 0.45; when also fitting *t*_0_, retinotopic: *t*_0_ = 209 ms, *R*^2^ = 0.75, non-ret.: *t*_0_ = 306 ms, R^2^ = 0.45). This finding suggests that perceptual learning does not mediate the correlation of RT_vert_ and TAE. The improved correlations compared to when measurements are averaged across days may reflect the larger variation in RTs (*SD* of 550 ms), or possibly that aggregating measurements across days having different RTs dilutes the observable effect of RT.

**Fig. 5.**
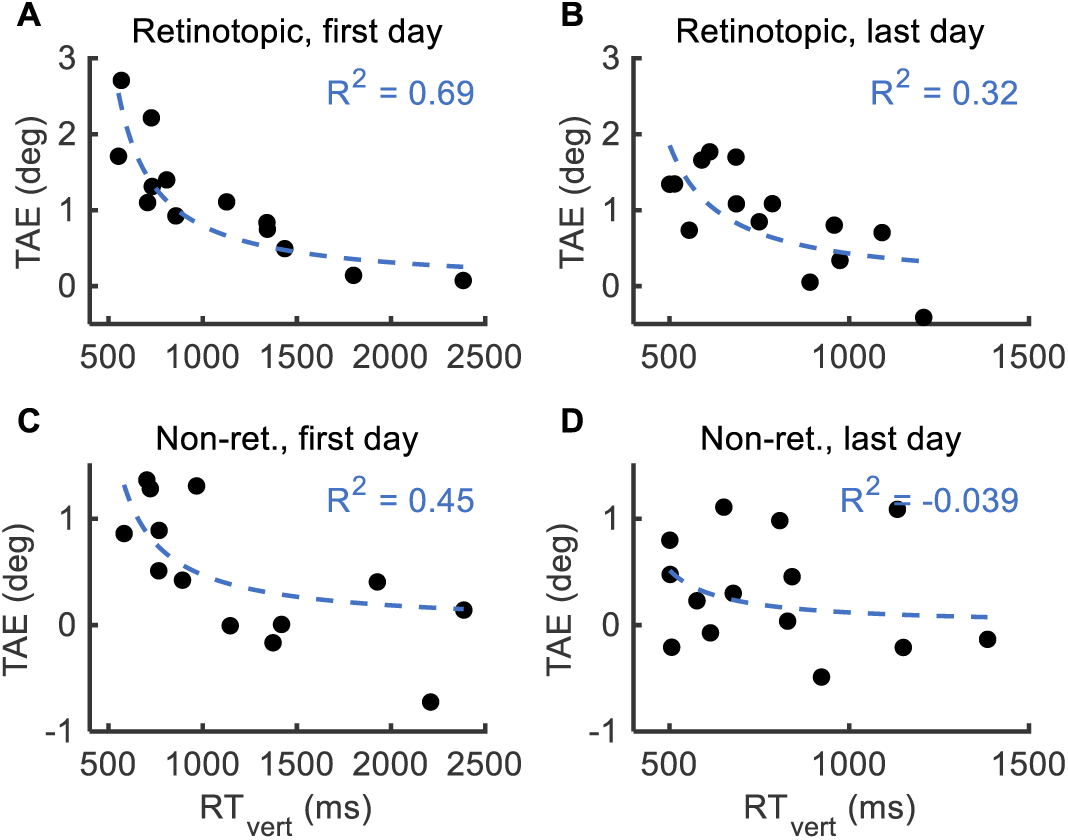
Co-variation of RT and TAE: the effects of experience with the task. Same as Fig. 4, for the (**A and C**) first and (**B and D**) last experimental sessions, of the (**A and B**) retinotopic and (**C and D**) non-retinotopic periphery experiment.

In the last session per observer (Fig. 5BD; day 5 ± 2, Mean ± *SD*), individual variation in RTs was significantly reduced compared with that on the first day (Mean ± *SD* of 770 ± 220 ms), a typical effect of practice (Harris & Sagi, 2018; Sagi, 2011); however, the correlation with TAE was still significant for the retinotopic TAE case (retinotopic: *R*^2^ = 0.6, *p* = 0.001, *t*_(12)_ = −4.24; non-ret.: *R*^2^ = 0.03, *p* = 0.5, *t*_(12)_ = −0.66; exclusion of the same observer as above leads to the same results; fitting to Eq. (19) showed, with *t*_0_ = 350 ms, retinotopic: R^2^ = 0.32, non-ret.: R^2^ = −0.03; when also fitting *t*_0_, retinotopic: *t*_0_ = 175 ms, R^2^ = 0.44; non-ret.: *t*_0_ = 377 ms, R^2^ = 0.02).

### Co-variation of RT and bias at fixation (TAE and TI)

First, we considered a TAE experiment, similar to the one above, in which all Gabor patches were presented at fixation (and hence retinotopic). The results showed a similar negative correlation between RT_vert_ and TAE; however, it was weaker than observed in the periphery (Fig. 6A) (*R*^2^ = 0.39, *p* = 0.03, *t*_(10)_ = −2.51; fitting to Eq. (19) with *t*_0_ = 350 ms: *R*^2^ = 0.16; when also fitting *t*_0_: *t*_0_ = 122 ms, *R*^2^ = 0.38). We also considered two tilt illusion experiments (TI, Fig. 1A, the spatial analogue of the TAE). The results showed a negative correlation between RT_vert_ and TI, though again the correlation was somewhat weak (Fig. 6BC) (“no jitter” experiment: *R*^2^ = 0.49, *p* = 0.03, *t*_(8)_ = −2.79; fitting to Eq. (19) with *t*_0_ = 250: *R*^2^ = 0.33; when also fitting *t*_0_: *t*_0_ = 0 ms, *R*^2^ = 0.45; “onset jitter” experiment: *R*^2^ = 0.47, *p* = 0.03, *t*_(8)_ = −2.66; fitting to Eq. (19) with *t*_0_ = 250: *R*^2^ = 0.15; when also fitting *t*_0_: *t*_0_ = 0 ms, *R*^2^ = 0.43). Overall, these findings replicate the observation for peripheral TAE, albeit at a smaller effect size. The weaker effect can be possibly explained by (i) reduced variation in RTs when tested at fixation (compare the x-axis in Fig. 4 to Fig. 6), (ii) less data per observer (3-8 daily sessions for TAE in the periphery, a single daily session for TAE in fixation and TI), or, most interestingly, (iii) the existence of a large RT-independent component in TAE and TI in these experiments (see within-individual analysis in Dekel & Sagi, 2019).

**Fig. 6.**
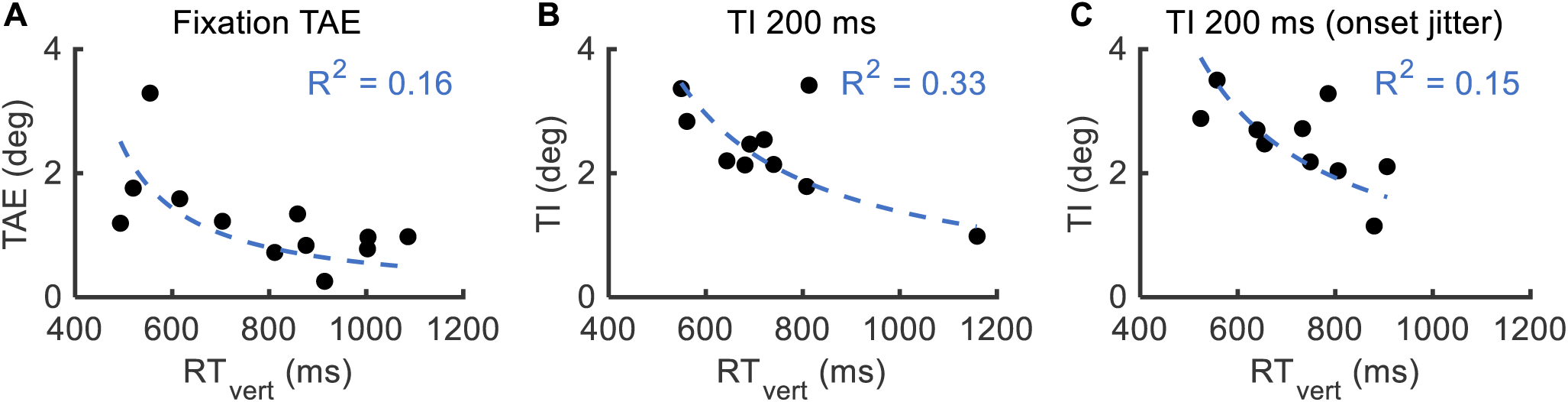
Co-variation of RT and bias in fixation (TAE and TI). The same as Fig. 4, for (**A**) the fixation TAE experiment, (**B**) the TI experiment with a 200 ms presentation duration and no onset jigger, and (**C**) the TI experiment with a 200 ms presentation duration and onset jigger. For the TI experiments, a fixed value of t_0_ = 250 ms was used in the fits. All experiments show a weak but significant correlation between bias magnitude and RT (p < 0.05).

### Just noticeable differences (JNDs)

Using the orientation discrimination task, it is possible to obtain a measure for orientation JND (just noticeable difference, calculated as a multiple of the interpolated inverse slope of the psychometric function; see the Methods). Considering JNDs is interesting given previous works suggesting that variability in TI can be explained by variability in JNDs (Song et al., 2013). Here, the results showed a relatively minor variation in JNDs between observers (*SD*/Mean of ~25%; Mean ± *SD* of JNDs across observers showing TAE retinotopic: 1.85° ± 0.58°, TAE non-retinotopic: 1.70° ± 0.42°, TAE fixation: 1.88° ± 0.47°, TI: 2.45° ± 0.47°; we noted that the small individual component was statistically significant in the TAE retinotopic condition with *p* = 1.2×10^−8^, *F*_(13,57)_ = 7.85). Given the small individual variability in JNDs, it is perhaps not surprising that we did not find any correlation between TAE or TI and JND (all *R*^2^ ≤ 0.1). Similarly, correlations between JND and RT_vert_ were not significant (TAE fixation: *R*^2^ = 0.31, *p* = 0.06, *t*_(10)_ = 2.10; TI “no jitter”: *R*^2^ = 0.23, *p* = 0.16, *t*_(8)_ = 1.53; all other conditions: *R*^2^ ≤ 0.05; correlations with 1/JND show the same). The lack of variation in JNDs may be attributed to the sampled population (see the Discussion).

### Are fast decisions caused by biased initial conditions?

As evident from changing *b* in Eq. (11), biased starting points inherently lead to faster decision times in the bounded decision model (DDM), even when the bound separation (*a*) is fixed. However, this property is unlikely to account for the correlation of bias and RTs observed here for the following reasons. (i) Fitting the retinotopic data to the DDM showed much better fits when *b* is fixed in the population, compared with when *a* is fixed in the population (log-likelihood of about −300 vs. −820; fits obtained using an exhaustive search over the *a* and *b* alternatives, and, for each alternative, finding the optimal drift rate and non-decision time using the fast-dm software, Voss & Voss, 2007). The same was found when using an unconstrained fit and correlating fitted *a* and *b* values with behavioral RT_vert_ (averaged within observer; correlation with *a*^*2*^: *R*^2^ = 0.88, *p* = 7×10^−7^, *t*_(12)_ = 9.39; correlation with *b*^2^: *R*^2^ = 0.01, *p* = 0.72, *t*_(12)_ = 0.36; again using fast-dm, but with a Kolmogorov-Smirnov setting and minimal outlier pruning). (ii) The retinotopic and non-retinotopic conditions exhibited different TAE magnitudes, but almost identical RTs (*R*^2^ = 0.99), suggesting a negligible influence of bias on RT. (Indeed, this strong independence can be taken as evidence in favor of an unbounded over a bounded decision process, consistent with Dekel & Sagi, 2019.) (iii) Conceptually, as seen in Eq. (11), the *b*^2^ term is subtracted from a large constant, so the range of possible RTs from changing *b* is more limited than from changing *a*.

Another noteworthy possibility for having different RTs in different individuals is that some observers have slower decision making, i.e., the scaling constant is changed between individuals (see *σ* in Eq. (15)). However, this alternative predicts that criterion shift bias (Eq. (1)) is invariant of time (see Eq. (12)), unlike behavior (e.g., the correlation of RT_vert_ and retinotopic criterion shift bias showing *R*^2^ = 0.62, *p* = 7.8×10^−4^, *t*_(12)_ = −4.46).

### Co-variation of retinotopic and non-retinotopic TAEs

In the experiments, the retinotopic and non-retinotopic trials differed only in their relative target position, leading to almost identical RTs in the two trial types (*R*^2^ = 0.99 in the population). Therefore, in the RT account of TAE described above, almost identical RTs were used in the retinotopic and the non-retinotopic cases (as evident in Fig. 3). Transitivity thus suggests a correlation between retinotopic and non-retinotopic TAEs, which was indeed found (Fig. 7) (*R*^2^ = 0.61, *p* = 0.001, *t*_(12)_ = 4.35). A similar, though non-significant correlation was found between the TI and the fixation TAE effect magnitudes (five shared observers, *R*^2^ = 0.45, *p* = 0.21, *t*_(3)_ = 1.58). This correlation is consistent with the idea of a common factor for similar visual phenomena (Grzeczkowski et al., 2017). Here, the common factor seems to be explained by RT, though a general account may also require other factors (such as JND, see the Discussion).

**Fig. 7.**
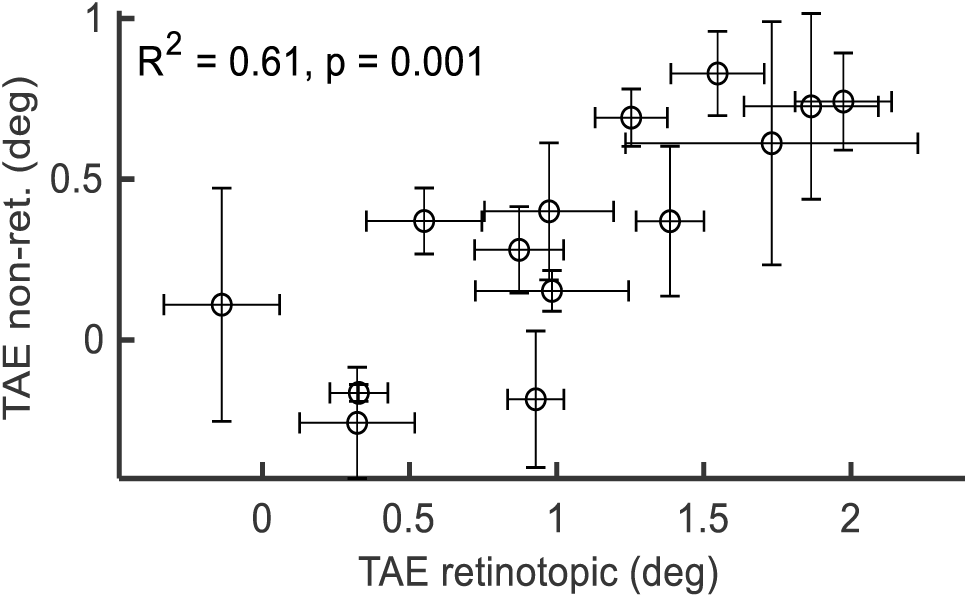
Co-variation of retinotopic and non-retinotopic TAEs. Non-retinotopic TAE is shown as a function of retinotopic TAE, for different observers, averaged across daily repetitions. Error bars are SEM across repetitions.

The strong correlation of non-retinotopic TAE with RT (Fig. 3) and with retinotopic TAE (Fig. 7) described above suggests that the non-retinotopic effect is not negligible, despite having a very small average magnitude (*M* = 0.33°, *p* = 0.005, *t*_(13)_ = 3.42, two-tailed t-test; this measurement is approximately consistent with previous reports, Knapen et al., 2010).

## Discussion

### TAE and RT

The results showed a large variability between individuals regarding their measured TAE magnitudes and high test-retest reliability. This variability is consistent with findings in earlier works for other context-dependent biases (Grzeczkowski et al., 2017; Song et al., 2013). The results also showed a large variation in RT between individuals, where RT ranged from ~500 ms to ~1800 ms when the target was vertical (Fig. 4). Importantly, the variability in TAE was strongly and negatively correlated with variability in RT (see Figs. 3–6). For example, in retinotopic TAE, the fastest observers had magnitudes of ~2°, whereas the slowest observers had almost no TAE. In the TI, we found a similar albeit weaker correlation with RT.

Based on modeling using evidence accumulation decision models, we concluded that the correlations suggest the involvement of decision confidence: to achieve a higher confidence level, decisions are slower, which we argue leads to reduced bias. Thus, we explain the individual differences by observers differing in their confidence level. Specifically, the RT-dependence of the TAE, evident between and within observers (Dekel & Sagi, 2019b), can be explained by the notion that TAE reflects a change in the initial conditions (priors) of a decision process. With decision time, the influence of the initial conditions is expected to be gradually reduced, leading to less bias in slower decisions. Note that even when no evidence exists (e.g., a vertical target orientation), we still expect the influence of the prior to be reduced with decision time (see the Methods, Dekel & Sagi, 2019). Note that here the time difference between the adaptation and test is always fixed; therefore, a slower RT does not correspond to increased adaptation decay (Greenlee & Magnussen, 1987; Magnussen & Johnsen, 1986). An alternative account, that highly biased observers are faster because their decision process started closer to a bound, was much worse in explaining the current data. (For the same reason, the possibility of individual differences in the rate of adaptation decay is not by itself sufficient to explain the current data.)

Overall, we found that individuals with a different bias can be explained by having different RTs but the same internal prior (see fits to Eq. (19) in Figs. 4-6; see the Methods; note the fixed *b* in modeling). This observation is important, both conceptually and technically, for understanding individual differences in vision, for example, when attempting to map the strength of measured aftereffects to internal visual priors (Grzeczkowski et al., 2017; Pellicano & Burr, 2012; Tibber et al., 2013; Yang et al., 2013). Behaviorally, we propose that RT-dependent bias is consistent with mechanisms that calibrate visual perception, for example, a prior for the reference frame of an object, which is gradually updated based on object details. (With spatial context, as in the tilt illusion, “prior” is reasonable in terms of coarse-to-fine processing.) Such calibration effects probably depend on probabilistic inference using the stimuli and are largely independent of neuronal constraints. For example, a change in gain or sensitivity due to contrast adaptation is probably time independent.

We noted that behavioral biases such as TAE and TI may reflect a sum over different underlying mechanisms (Bao & Engel, 2012; Dekel & Sagi, 2019a). Possibly not all bias mechanisms are RT dependent. An RT-independent bias component can be modeled using evidence accumulation theories as a context-dependent change in the drift rate (Dekel & Sagi, 2019b; Summerfield & De Lange, 2014). As such, the analysis described here needs to be adjusted in order to apply to possible experimental conditions where an RT-independent component dominates the measured bias.

### TAE and JND

There seems to be strong evidence that context-dependent biases are positively correlated with (measured) orientation JND, both within-observer (Solomon & Morgan, 2006; Wei & Stocker, 2017), and between observers (TI: Song et al., 2013, color and face aftereffects: Mattar, Carter, Zebrowitz, Thompson-Schill, & Aguirre, 2018). The JND-dependence of bias seems consistent with a type of bias that depends on the constraints of the system, such as the bandwidth of orientation-selective units or the size of V1 (Song et al., 2013). This idea is not incompatible with the mechanism proposed above, because measured TAEs may correspond to a sum of effects from multiple mechanisms having different objectives and properties (see Bao & Engel, 2012). Interestingly, we find that JND-dependent bias is predicted in decision models from individual differences in the duration (Eqs. (10) and (17)) and the rate (Eq. (18)) of evidence accumulation.

Here, the results showed no correlation between TAE or TI and JND. The lack of a strong correlation can be explained by the small variation in JNDs in the sampled population (*SD/*Mean of ~25%, much less than in Song et al., 2013). The lack of variation in JNDs may be attributed to measuring JNDs for perceived orientation (no physical reference), to rapid saturation of perceptual learning within the first session, or to the sampled population, which consisted of observers from the same age group (18-40) with comparable visual acuity (6/6). (We wish to emphasize that from a reductionist perspective, we found it important to understand individual differences while controlling for visual acuity and age.) Generally we expect a full account of individual variation in context-dependent bias to depend on both RT and JND.

Based on both bounded and unbounded models (see Eqs. (10) and (17)), sensitivity is expected to increase in proportion to the square-root of the decision time. However, such a dependence was not found in the behavioral data. This may be explained by the smaller size of the expected effect (the square-root of the size of the effect of RT vs. TAE). Another possibility is of a discrepancy between the assumption using both models that the evidence is fixed in time, and the experiments, where the evidence was temporary (a target presentation duration of 50 ms with possibly a few hundred more milliseconds of persistence). This observation warrants caution when using simple models to interpret behavior. Still, for the purpose of this work, the analysis of RT and bias seems robust for a dynamic reduction in the evidence, because the found dependence of bias on RT persists in the case where there is no evidence (i.e., *v* = 0; see the Methods).

Note that if bias magnitude is only explained by the intrinsic orientation sensitivity of the system (i.e., having large bias when accuracy is low, Song et al., 2013), and the less sensitive observers compensate by responding more slowly (thus, measured JND is fixed, and measured RT is variable), then we expect a positive correlation between bias and RT, the opposite of what was observed here. Therefore, even assuming a speed-accuracy tradeoff, our results cannot be explained by the existing work (Song et al., 2013). (Possibly, antagonistic correlations of JND and TAE, both positive and negative, led to the net zero correlation found here.)

### Spatially non-selective TAE

Previous work suggested that the average TAE is very weak in non-retinotopic settings (i.e., when the adaptor and the target are presented at different retinal positions) (Knapen et al., 2010). We replicated this finding, but importantly, we found that the non-retinotopic TAE was strongly correlated with both RT (Fig. 3) and retinotopic TAE (Fig. 7). These findings suggest that the effect is more important than previously thought.

## Acknowledgments

This research was supported by the Basic Research Foundation, administered by the Israel Academy of Science, and by The Weizmann Braginsky Center for the Interface between the Sciences and the Humanities. We thank Dr. Misha Katkov, Dr. Noga Pinchuk-Yacobi, and Michelangelo Naim for their suggestions and comments.

